# Generation of a transparent killifish line through multiplex CRISPR/Cas9-mediated gene inactivation

**DOI:** 10.1101/2022.07.04.498720

**Authors:** Johannes Krug, Carolin Albertz, Vera L. Hopfenmüller, Christoph Englert

## Abstract

Body pigmentation is a major limitation for *in vivo* imaging and thus for the performance of longitudinal studies in biomedicine. A possibility to circumvent this obstacle is the employment of pigmentation mutants, which are used in fish species like zebrafish and medaka. To address the molecular basis of aging, the short-lived African killifish *Nothobranchius furzeri* has recently been established as a model organism. Despite its short lifespan, *N. furzeri* shows typical signs of mammalian aging including telomere shortening, accumulation of senescent cells and loss of regenerative capacity. Here, we report the generation of a transparent *N. furzeri* line by simultaneous inactivation of three key loci responsible for pigmentation. We demonstrate that this stable line, named *klara*, can serve as a tool for different *in vivo* applications including behavioral experiments addressing mate choice and the establishment of a senescence reporter by homology-directed repair-mediated integration of a fluorophore into the *cdkn1a (p21)* locus.

## Introduction

In animals, pigments that can be found in specific cell types limit optical transparency and prevent the *in vivo* observation of processes like organogenesis, regeneration or cancer metastasis. While mammals have only one pigment cell type, the melanocyte, other vertebrates including fish develop several chromatophores that produce different colors. In one of the best-studied models for vertebrate coloration, the zebrafish (*Danio rerio*), the three main kinds of chromatophores are the melanophores (black), the iridophores (silvery or blue) and the xanthophores (yellow), all derived from neural crest cells ^1,2^. A fourth population of pigment cells, forming the retinal pigment epithelium (RPE) is derived from the optic neuroepithelium ^3^. Different combinations of naturally occurring mutants in pigmentation genes have been used to generate adult transparent zebrafish. The *casper* line lacks melanocytes and iridophores due to mutations in *mitfa* and *mpv17*, respectively ^4,5^. An additional mutation in the *slc45a2* gene is present in *crystal* zebrafish, which completely lack melanin and therefore possess a transparent RPE ^6^. Transparent zebrafish have been used to study different aspects of cancer and stem cell biology, among others ^5,7^. In another model fish, the medaka (*Oryzias latipes*), transparent juvenile and adult animals have recently been generated through CRISPR/Cas9-mediated inactivation of *oca2* and *pnp4a* ^8^.

During the last decade the turquoise killifish, *Nothobranchius furzeri*, has emerged as a new model for research on aging ^9^. With a lifespan between three and seven months, *N. furzeri* is the shortest-lived vertebrate that can be kept in captivity ^10,11^. Hatchlings grow rapidly and can reach sexual maturation already within two to three weeks ^12^. *N. furzeri* shares many hallmarks of aging with mammals, including telomere shortening, mitochondrial dysfunction, cellular senescence, loss of regenerative capacity and cognitive decline ^13–19^. Despite its short lifespan, *N. furzeri* also shows a high incidence of age-dependent neoplasias in liver and kidney ^20^. What makes the killifish an attractive model in addition, is the establishment of transgenesis and genome engineering ^15,21–23^ as well as the availability of reference sequences for the *N. furzeri* genome ^24,25^. The turquoise killifish is sexually dimorphic and dichromatic ^26^. Compared to females, males are larger and colorful. The latter occur in two color forms with red and yellow morphs that differ primarily in coloration of the caudal fin. Females have translucent fins and a pale greyish body with iridescent scales.

Here, we describe the generation of fully transparent juvenile and adult *N. furzeri* animals. We have used a single injection of three sgRNAs targeting *mitfa, ltk* and *csf1ra*, which are involved in pigment development in melanophores, iridophores and xanthophores, respectively. With the method employed we have achieved simultaneous and biallelic somatic gene disruptions of three genomic loci in a highly efficient manner. Already in the F_0_ generation, a fraction of animals were fully transparent. Homozygous triple mutants showed normal behavior, fertility and general health. In addition, we have used the transparent line, named *klara*, to inactivate additional genes, to study female and male mate choice and to generate an *in vivo* senescence reporter by homology-directed repair-mediated integration of an *GFP* allele into the locus of the senescence marker *cdkn1a (p21)*.

## Results

### Multiple genes can be simultaneously inactivated in *N. furzeri*

For the generation of a transparent *Nothobranchius furzeri* line, we selected the genes *mitfa, ltk*, and *csf1ra* as targets to interfere with the formation of melanophores, iridophores and xanthophores, respectively. The expression of those three genes was analyzed in skin tissue from fish of both sexes at the age of 1, 2, 3 and 6 weeks post hatching (wph). In male fish, we observed a significant up-regulation of *mitfa* at the age of 6 wph, whereas *mitfa* expression did not change in female fish (Fig. 1a). The expression of *ltk* increased steadily with age in both, male and female fish (Fig. 1b). In contrast, *csf1ra* expression did not differ significantly over time in males and females (Fig. 1c). Direct comparison of *mitfa, ltk* and *csf1ra* expression between females and males did not reveal sex-specific differences except for *mitfa* at 6 wph (Extended Data Fig. 1a-c). To induce mutations in the selected genes, single guide RNAs were designed based on the genome sequence provided by the *Nothobranchius furzeri* Genome Browser ^24^. Since three genes should be inactivated at the same time, we used one sgRNA per gene. To facilitate mutation detection PAM sequences were chosen that had a restriction site directly upstream, assuming that the restriction site would be lost upon the introduction of a mutation. After characterization of different sgRNAs for each gene, sgRNAs were synthesized targeting *mitfa* in exon 6, *ltk* in exon 22 and *csf1ra* in exon 9 (Extended Data Fig. 1d-f). To simultaneously inactivate *mitfa, ltk* and *csf1ra*, we injected *Cas9* mRNA, the three different sgRNAs and *GFP* mRNA into one-cell-stage embryos of the long-lived *N. furzeri* strain MZCS-08/122 ^27^. GFP mRNA was used to indicate properly injected embryos one day after the injection. An injection mold, stabilizing the eggs during the injection procedure, has already been reported ^28^. To further improve the injection procedure, we developed a new type of injection mold having single slots for each embryo. Moreover, the wall of these slots pointing towards the direction of the injection needle is sloped facilitating the access of the needle to the egg (Extended Data Fig. 1g). Using this injection mold, we were able to inject close to 600 embryos within two days, which were sorted for a GFP signal one day post injection. One third each of the embryos were GFP-positive, GFP-negative or dead (Extended Data Fig. 1h). Seven days after injection, GFP-positive eggs were transferred onto coconut coir plates, mimicking the dry phase *N. furzeri* eggs undergo in their natural habitat. Since it is possible to detect melanophores already in embryos, GFP-positive and GFP-negative embryos were phenotypically analyzed. While melanophores were clearly detected in GFP-negative embryos, a reduction or an almost complete loss of melanophores was observed on the head and along the body axis in a fraction of GFP-positive embryos (Fig. 1d). Iridophores and xanthophores are not detectable at this stage of development.

**Fig.1.**
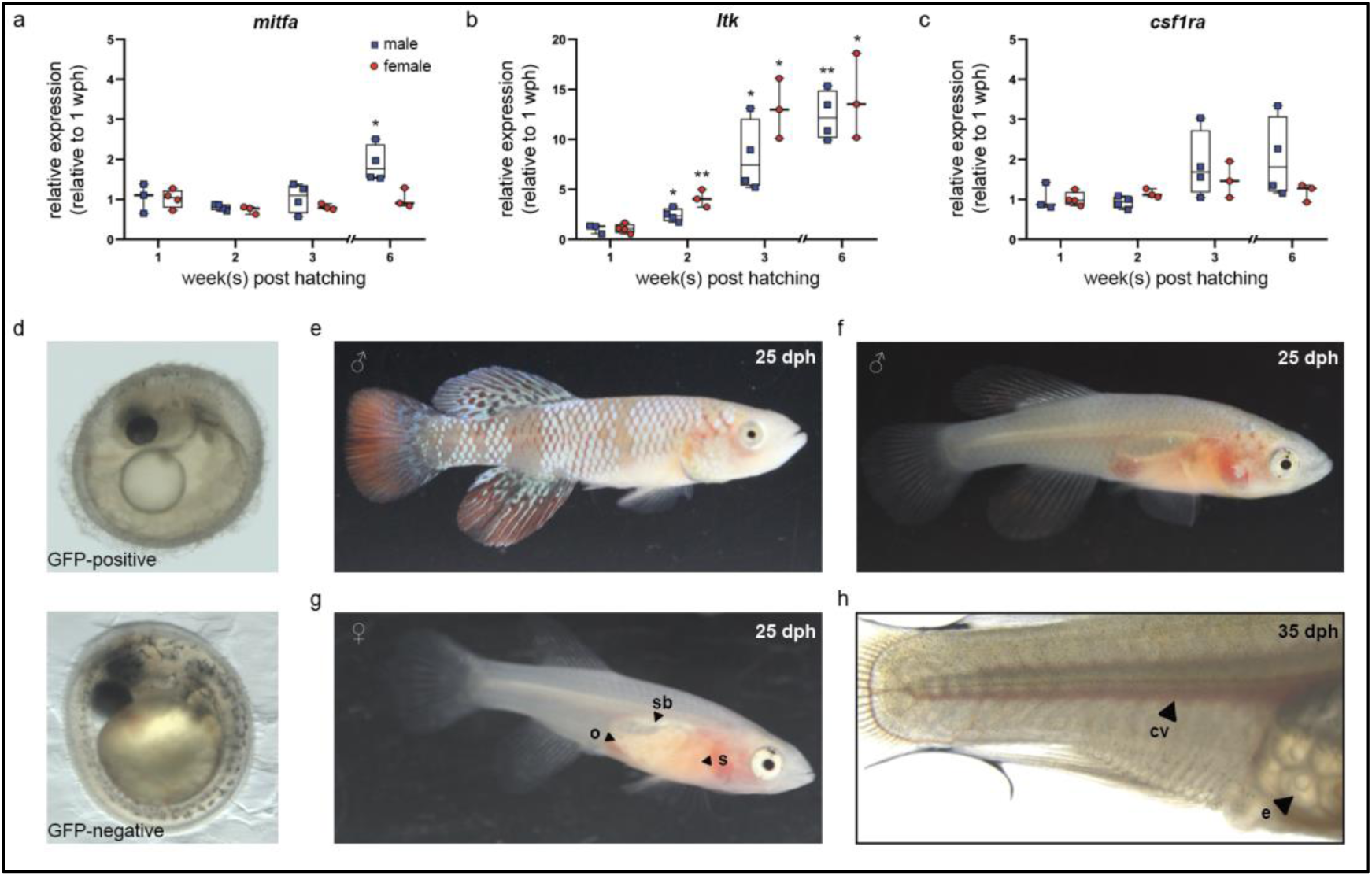
Simultaneous inactivation of genes involved in body pigmentation. **a**,**b**,**c**, Gene expression analysis of *mitfa* **(a)**, *ltk* **(b)** and *csf1ra* **(c)** in the skin of male and female wild type fish via quantitative real-time PCR. The expression of *mitfa* was only significantly up-regulated in male fish at the age of 6 weeks post hatching (wph) (p: 0.048). For both sexes, the expression of *ltk* increased significantly over time (p_male, 2wph_: 0.028, p_male, 3wph_: 0.026, p_male, 6wph_: 0.002; p_female, 2wph_: 0.002, p_female, 3wph_: 0.019, p_female, 6wph_: 0.033), whereas no differences were observed for *csf1ra*. Expression levels were normalized to the expression at 1 wph in the respective sex. *rpl13a* was used as housekeeping gene. (n_male_1wph_= 3, n_male_2wph_= 4, n_male_3wph_= 4, n_male_6wph_= 4; n_female_1wph_= 3, n_female_2wph_= 4, n_female_3wph_= 4, n_female_6wph_= 4). Relative gene expression was calculated using the ΔΔCT method. Student’s or Welch’s t-tests were computed to determine significant changes in gene expression. **d**, Phenotypical analysis of embryos revealed a reduction of melanophores in GFP-positive compared to GFP-negative embryos. **e**, In fish of the F_0_ generation, a mosaic loss of body pigmentation was observed. **f**,**g**, In addition, almost fully transparent F_0_ fish were detected allowing a view on inner organs (o: ovary, s: stomach, sb: swim bladder). **h**, Microscopic analysis of a female F_0_ fish with a view on individual eggs within the ovary and blood vessels (cv: cardinal vein, e: egg).

In order to analyze if the sgRNAs targeting *mitfa, ltk* and *csf1ra* had induced mutations, regions around the expected mutation sites were amplified via PCR using DNA extracts from 8 randomly selected GFP-positive and 6 GFP-negative embryos. Those amplicons were used in restriction enzyme digests to determine the presence of mutations. For all three genes. we observed a non-cleaved PCR fragment in all samples of GFP-positive embryos, suggesting that mutations had been introduced that made the amplicons resistant to restriction enzyme digest. The analysis of mutations in the *mitfa* sequence revealed, that 75% (6/8) of the GFP-positive samples only showed one undigested fragment, whereas 25% (2/8) were mosaic, since they also showed two additional fragments that only occur in the presence of the wild type sequence. For *ltk*, we observed mosaicism in 37.5% of embryos (3/8) and for *csf1ra* in 87.5% (7/8) (Extended Data Fig. 1i-k). For the GFP-negative embryos, we identified digested fragments in all samples, as in the respective wild type controls. However, the first two GFP-negative embryos also showed non-cleaved fragments, indicating the presence of mutations. This suggests that the sorting into GFP-positive and GFP-negative embryos is not fully stringent.

Next, we hatched and raised GFP-positive embryos from the F_0_ generation. Compared to wild type larvae and adult animals, the majority of fish from the F_0_ generation of successfully injected eggs showed a mosaic loss of pigment cells. However, in some of the fish we could already observe an almost complete loss of pigment cells, resulting in transparency of the animals (Fig. 1e-h). This already allowed us to have a clear view on inner organs. At an age of 25 days post hatching (dph), the stomach in those transparent fish shows an orange color due to artemia, small crustaceans, which are used as food at this age. Moreover, the swim bladder and in females the ovaries were clearly visible. One could also observe the blood flow in the cardinal vein and in the small vessels of the caudal fin. Additionally, we could identify single eggs including lipid droplets in the ovaries. Besides this phenotypical analysis of fish from the F_0_ generation, we also analyzed the mutation rates. Based on results from the previously mentioned restriction analysis, we observed that all 85 fish had a mutation in *mitfa, ltk* and *csf1ra*, at least in a mosaic fashion. Moreover, this analysis revealed that already in the F_0_ generation biallelic mutations were observed (*mitfa*: 48.2%, *ltk*: 67.1%, *csf1ra*: 23.5%) (Extended Data Fig. 1l). These data indicate a high efficiency of the CRISPR/Cas9 tool in *N. furzeri* and show that it is possible to simultaneously induce mutations in three genes of interest.

### Generation of a stable, transparent killifish line

For the generation of a stable, transparent *N. furzeri* line and to reduce potential off-target effects, we performed an outcross of selected F_0_ fish with wild type animals. As expected, all of the obtained F_1_ progeny showed a normal pigmentation pattern and hence were phenotypically not distinguishable from wild type animals. To assess the presence of mutations, we extracted DNA from fin biopsies for genotyping. Among 60 analyzed fish, 14 animals were triple-heterozygous (*mitfa*^*+/−*^, *ltk*^*+/−*^, *csf1ra*^*+/−*^). The targeted loci in those fish were analyzed via sequencing. Various deletion mutations were detected, but two fish carried the same mutation at the *mitfa* (Δ11 bp), *ltk* (Δ4 bp) and *csf1ra* (Δ5 bp) locus (Extended Data Fig. 2a-c). Those animals were used for a subsequent incross, from which according to Mendelian ratio 1/64 (1.56%) of the F_2_ offspring was expected to be triple homozygous. We first checked whether triple homozygous embryos were viable and could be detected among the F_2_ eggs. Hence, we randomly selected embryos to assess their genotype. We observed a lack of melanophores in a proportion of embryos. Those embryos carried a homozygous mutation in *mitfa*, whereas embryos with only a heterozygous *mitfa* mutation had melanophores (Extended Data Fig. 2d-k). The genotypes for *ltk* and *csf1ra* could only be assessed via molecular analysis at this developmental stage. Notably, one triple homozygous embryo was detected, indicating that at least until the hatching stage those embryos were viable. We then proceeded with hatching of F_2_ eggs. In this generation, different genotypes have been observed to result in different phenotypical appearances. Female fish with a heterozygous mutation in *ltk* and homozygous mutations in *mitfa* and *csf1ra* resembled at first glance wild type females, whereas in male fish especially the lack of xanthophores resulted in a silver-blue appearance (Fig. 2a,a’). In contrast to this, the lack of melanophores and iridophores in fish with the genotype *mitfa*^*-/-*^, *ltk*^*-/-*^, *csf1ra*^*+/−*^ allowed a view on inner organs, particularly in females (Fig. 2b,b’). In order to increase the likelihood to obtain transparent, triple homozygous fish, we crossed two fish with the genotype *mitfa*^*-/-*^, *ltk*^*-/-*^, *csf1ra*^*+/−*^. From this cross we genotyped 50 individuals via high-resolution melting analysis (HRMA) and could identify 13 triple homozygous fish. We named the transparent *N. furzeri* line *klara* (Fig. 2c,c’).

**Fig.2.**
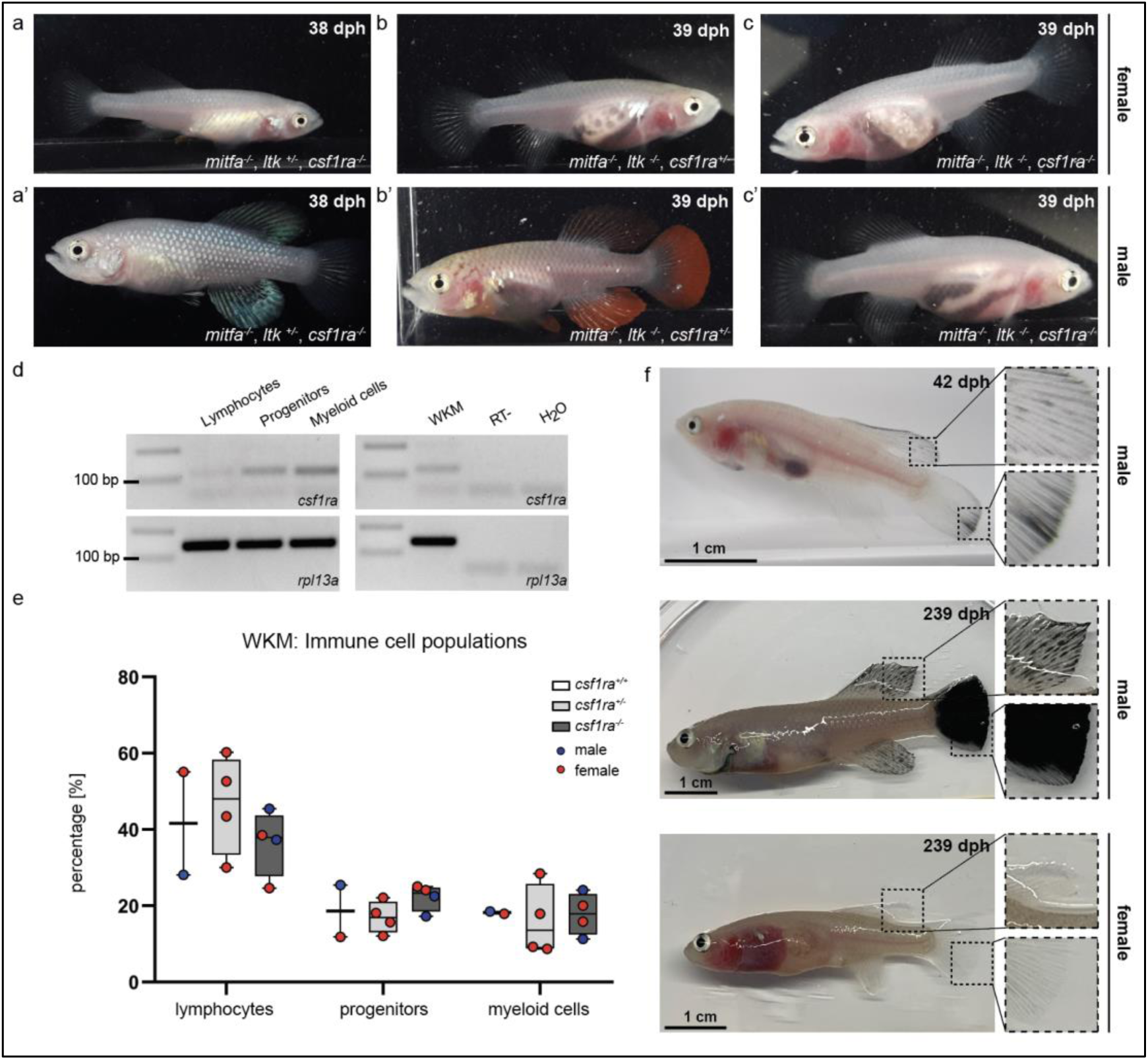
Generation and characterization of *klara*. **a**,**a’**, Female **(a)** and male **(a’)** fish at the age of 38 days post hatching (dph) with the genotype *mitfa*^*-/-*^,*ltk*^*+/−*^,*csf1ra*^*-/-*^ showed a lack of melanophores and xanthophores, whereas iridophores were present. **b**,**b’**, A lack of melanophores and iridophores was observed in female **(b)** and male **(b’)** *N. furzeri* with the genotype *mitfa*^*-/-*^,*ltk*^*-/-*^,*csf1ra*^*+/−*^. Despite a homozygous mutation in *ltk* (*ltk*^*-/-*^) individual scales with iridophores were detected in fish of both sexes. **c**,**c’**, The presence of homozygous mutation in all the three genes *mitfa, ltk* and *csf1ra* resulted in a loss of body pigmentation in females **(c)** and males **(c’)** allowing a view on inner organs. **d**, Expression of *csf1ra* was analyzed via RT-PCR using cDNA from FACS-sorted populations of lymphocytes, progenitors and myeloid cells obtained from the whole kidney marrow (WKM) of a wild type *N. furzeri. Csf1ra* was detected in all subpopulations, most strongly in myeloid cells. As negative control, an RT-sample (no reverse transcriptase during cDNA synthesis) was used to exclude contaminations with genomic DNA. As loading control, *rpl13a* was used. **e**, Comparison of cell numbers in the different subpopulations of the WKM of fish with the following genotypes: *mitfa*^*-/-*^,*ltk*^*-/-*^, *csf1ra*^*+/+*^ (n= 2), *mitfa*^*-/-*^,*ltk*^*-/-*^,*csf1ra*^*+/−*^ (n= 4) and *mitfa*^*-/-*^,*ltk*^*-/-*^,*csf1ra*^*-/-*^ (n= 4). One-way ANOVA followed by Tukey’s post hoc test did not reveal any significant differences. **f**, Male *klara* fish showed an appearance of melanophores on fin appendages, which intensified with age resulting in black fins. In female *klara* animals black fins were not observed.

### Characterization of *klara* animals

In zebrafish, the role of *csf1ra* in xanthophore development has been described ^29^. In addition, *csf1ra* is also known to play a role in the immune system, in particular in the survival, proliferation and differentiation of monocytes and macrophage ^30,31^. For this reason, we wondered whether the inactivation of *csf1ra* has an effect on the immune cell population of *klara*. Since the kidney is the primary hematopoietic organ in teleost fish, we analyzed the whole kidney marrow (WKM) of fish with homozygous mutations in *mitfa* and *ltk*, which had in addition either no mutation, a heterozygous or a homozygous mutation in *csf1ra*. Based on a published gating strategy ^32^, we could identify four subpopulations in the WKM of *N. furzeri* using flow cytometry (Extended Data Fig. 2l). The strongest *csf1ra* expression was detected in the myeloid cell population, which contains macrophages (Fig. 2d). Comparing the number of immune cells in the three sub-populations, we did not observe any differences among the *csf1ra* genotypes (Fig. 2e and Extended Data Fig. 2m).

During raising of *klara* fish, we observed that around the age of approximately four weeks, i.e., at the time of sexual maturation, melanophores appeared in male fish, in particular on fin appendages. However, this was not detected in female *klara* fish (Fig. 2f). With age the melanophores spread over the whole fins and resulted e.g., in a fully black caudal fin. This, however, did not interfere with the overall transparency of *klara* animals.

### Males and females prefer pigmented mating partners

Besides its role as camouflage, protection from UV damage or for recognition, body pigmentation also plays an important role for the choice of mating partners. We wanted to investigate whether the lack of body pigmentation affects the breeding behavior of *klara* fish. We set up three breeding groups consisting of one *klara* male and two *klara* females, at the age of 14 weeks, which had never been used for breeding before. *Klara* fish showed a normal mating behavior, whereby the male uses its caudal fin to push the female into the sand and thus induces egg laying (Extended Data Movies 1,2). We also assessed the quantity and quality of collected eggs, which did not differ from wild type fish, so that we could maintain the *klara* line in a triple homozygous state (Fig. 3a). To assess the role of body pigmentation for mate choice in killifish, we set up different combinations of breeding trios consisting of wild type and *klara* fish, thus that a wild type or a *klara* animal of each sex had the choice between a wild type or a *klara* animal of the other sex (Extended data Fig. 3). For each of the four combinations, we analyzed three tanks. Fish were put together and a sand box, which is required for egg deposition, was put into the tank. After 10 days the sand box was removed for the following two days. Subsequently, the box was added again and for the following four weeks we collected eggs once per week for further analysis. In order to decide who produced the egg or who had fertilized it, we determined the genotype of the fertilized eggs by HRMA. We observed that in the presence of a *klara* and a wild type female fish, irrespective of whether the male was a wild type or a *klara* animal, approximately 75% of fertilized eggs originated from the wild type female (Fig. 3b). This indicated that both *klara* and wild type males showed a preference for the pigmented wild type female. This mate choice was not influenced by size or weight, since both parameters were indistinguishable between wild type and *klara* females (Fig. 3c). Similarly, in the breeding groups, in which a *klara* and a wild type male was present, more than 90% of eggs were fertilized by the wild type male (Fig. 3b). Again, wild type and *klara* males did not show a difference in size, although *klara* males had less weight (Fig. 3c). Taken together, this competitive breeding experiments indicated that pigmented fish were the preferred mating partner for both sexes.

**Fig.3.**
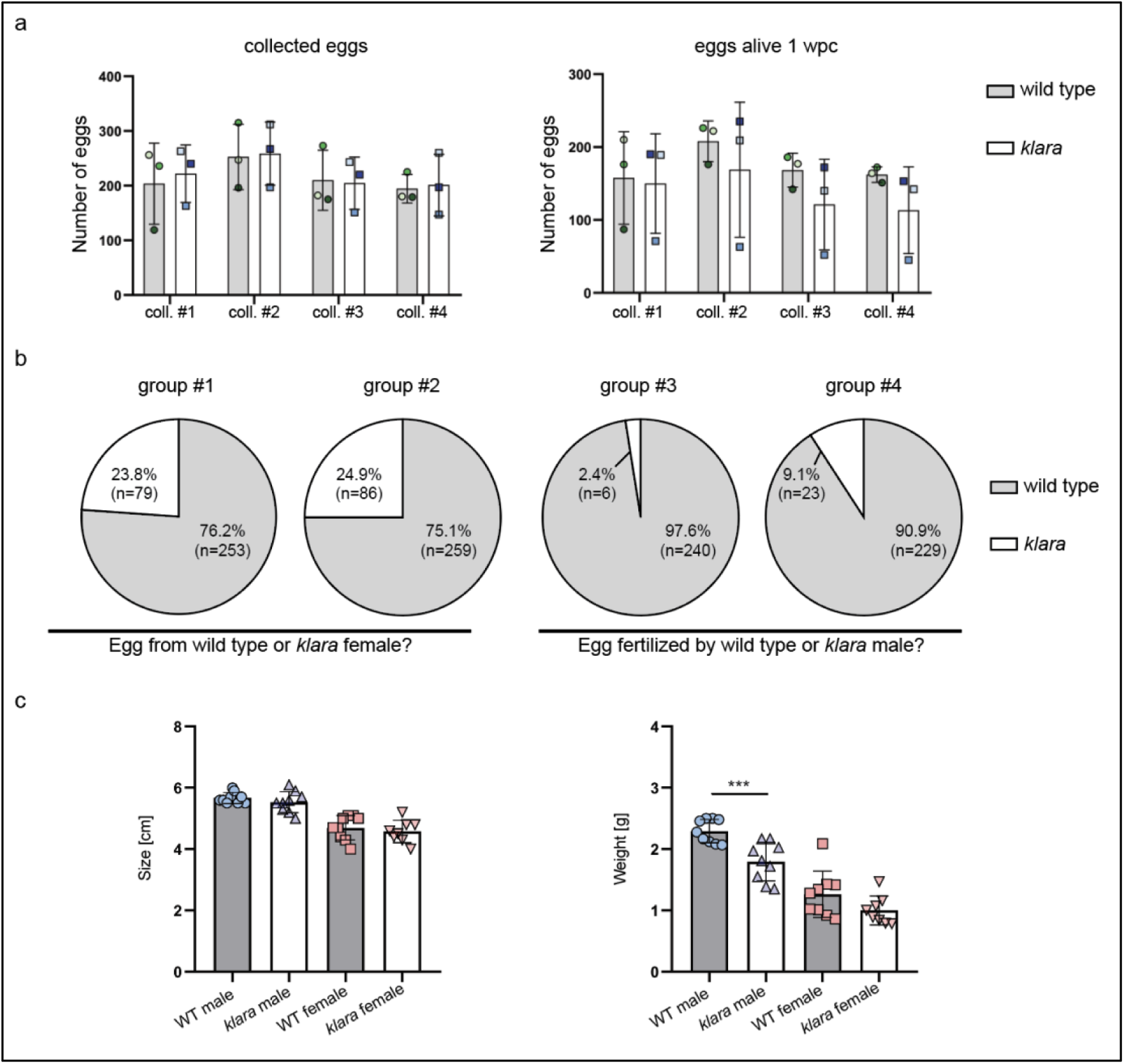
Pigmented fish are preferred mating partners. **a**, For the analysis of egg quantity, eggs were collected once per week for four weeks from wild type (n= 3 tanks with 1 male and 2 females each) and *klara* (n= 3 tanks with 1 male and 2 females each) fish. To asses egg quality, the amount of alive eggs stored on coconut coir plates was determined 1 week post collection (wpc). No significant differences were observed regarding egg quantity and quality. **b**, Collected eggs obtained from breeding groups with a wild type female and a *klara* female and either a wild type male (group #1) or a *klara* male (group #2) were genotyped via HRMA. As mating partner, male fish preferred wild type females (group #1: 76.2%; group #2: 75.1%) over *klara* females (group #1: 23.8%; group #2: 24.9%). In the presence of a wild type male and a *klara* male and either a wild type female (group #3) or a *klara* female (group #4) the majority of analyzed eggs were fertilized by the wild type male (group #3: 97.6%; group #4: 90.9%). **c**, Within the same sex, wild type and *klara* fish did not differ in size(n_WT_male_= 9 n_WT_female_= 9, n_*klara*_male_= 9, n_*klara*_female_= 8). Wild type males were significantly heavier than *klara* males (p: 0.0009). Student’s or Welch’s t-tests were computed to determine differences in size or weight.

### *Klara* fish can serve as a platform for further genetic manipulation

The simultaneous inactivation of *mitfa, ltk* and *csf1ra* resulted in a loss of body pigmentation, however, the eyes were still normally pigmented (Fig. 4a). Transparent zebrafish from the *casper* line ^5^ also still have pigmented eyes, while fish of the fully transparent *crystal* line lack those pigments. This was achieved via an inactivation of the *slc45a2* gene ^6^. In order to get rid of retinal pigmentation in *klara* animals we designed a sgRNA targeting the *slc45a2* locus (Extended Data Fig. 4a). We first tested this sgRNA in one-cell stage eggs from the wild type strain. Compared to GFP-negative embryos, we observed a loss of pigmentation in the eye of GFP-positive embryos (Fig. 4b). Notably, also the appearance of melanophores on the head and body of those embryos was reduced. This phenotypical analysis indicated inactivation of *slc45a2*, which was subsequently confirmed via restriction enzyme digest (Extended Data Fig. 4b). We then performed microinjections using the sgRNA targeting *slc45a2* into *klara* one-cell stage embryos and hatched F_0_ fish (n=7). Besides the transparent body, the black pigmentation of the eye was absent, while we still observed a silver-pigmented ring around the eye of the animals (Fig. 4c). The presence of a mutation in the *slc45a2* locus of all seven F_0_ fish was confirmed via restriction enzyme digest (Extended Data Fig. 4c). Subsequently, we performed an outcross of F_0_ fish with *klara* and identified the presence of various indel mutations in F_1_ offspring (Extended Data Fig. 4d,e). This experiment showed that the *klara* line can serve as a platform for further genetic manipulations.

**Fig.4.**
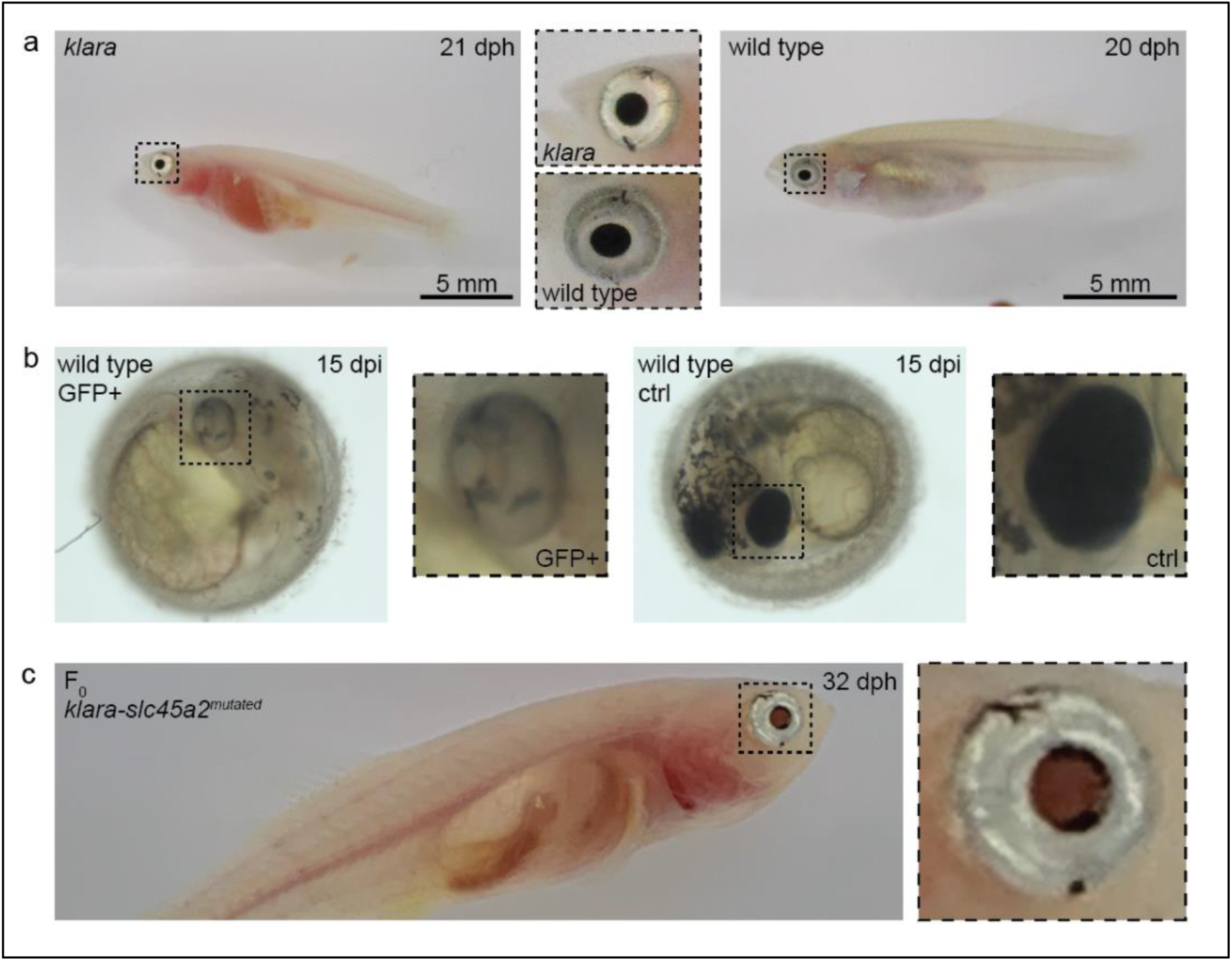
Loss of eye pigmentation. **a**, Despite the inactivation of three genes involved in pigmentation, the eye pigmentation did not differ between *klara* and wild type fish. **b**, Targeting the *slc45a2* locus in wild types, resulted in a reduction of melanophores in the eye, but also on the whole body of injected embryos compared to wild type controls. **c**, A loss of melanophores in the retinal pigmented epithelium was observed in F_0_ fish upon a microinjection of the sgRNA targeting *slc45a2* in *klara* embryos. A silver-pigmented ring around the eye was still present. (dpi: days post injection; dph: days post hatching)

### Generation of an *in vivo* senescence reporter

The accumulation of senescent cells is one of the hallmarks of the aging process ^33^. We wanted to take advantage of the transparent *klara* line and generate a senescence reporter line. To this end, we planned to insert a reporter construct, consisting of an eGFP and nitroreductase (NTR) cassette into the *cdkn1a* (*p21*) locus of *klara* via homology-directed repair (Fig. 5a). The *cdkna1* gene is a senescence marker and upregulated in old killifish ^14^. With this reporter/NTR cassette, *cdkn1a* expressing cells can be labelled via eGFP and can also be ablated via the NTR/Mtz system ^34^. To facilitate HDR, we added flanking arms of 903 bp and 901 bp on the 5’ and 3’ ends of the construct, respectively. Since it has been reported that 5’ modifications of double-stranded donor templates increase HDR efficiency ^35,36^, we amplified the template using biotinylated oligonucleotides (Fig. 5b). Subsequently, we performed microinjections into one-cell stage *klara* eggs using an injection solution with a sgRNA that induces a DNA double-strand break in close proximity to the intended insertion site and the biotinylated HDR donor template. In 1 out of 16 randomly selected GFP-positive (successfully injected) embryos, we detected the presence of the reporter cassette. Its proper insertion was verified via sequencing (Extended Data Fig. 5a). We subsequently set up the remaining GFP-positive embryos for hatching and detected three fish with a proper insertion among 19 F_0_ animals. Next, we performed an outcross of the F_0_ animals with *klara* fish. To assess whether the reporter was functional, the obtained F_1_ embryos were exposed to γ–irradiation of 10 Gy. This should induce DNA damage and lead to activation of TP53 and up-regulation of *cdkn1a*. To analyze, whether the up-regulated *cdkn1a* expression would also be linked to an up-regulation of *GFP*, we analyzed the embryos 1h before as well as 24h post irradiation (hpi) via fluorescence microscopy (Extended Data Fig. 5b). Before irradiation, autofluorescence originating from the yolk was observed in *cdkn1a*^*+/+*^ and *cdkn1a*^*ki/+*^ embryos (Fig. 5c). In contrast, at 24 hpi we observed GFP-positive cells in the optic tectum and other parts of the developing brain and fin buds of *cdkn1a*^*ki/+*^ embryos (Fig. 5d). Gene expression analysis revealed an upregulation of endogenous *cdkn1a* as well as both *GFP* and *ntr* expression upon γ–irradiation in *cdkn1a*^*ki/+*^ embryos, which further significantly increased at 48 hpi (Fig. 5e-g). Thus, we were able to insert a functional 1.5 kb reporter cassette into the *cdkn1a* locus of *klara* fish.

**Fig.5.**
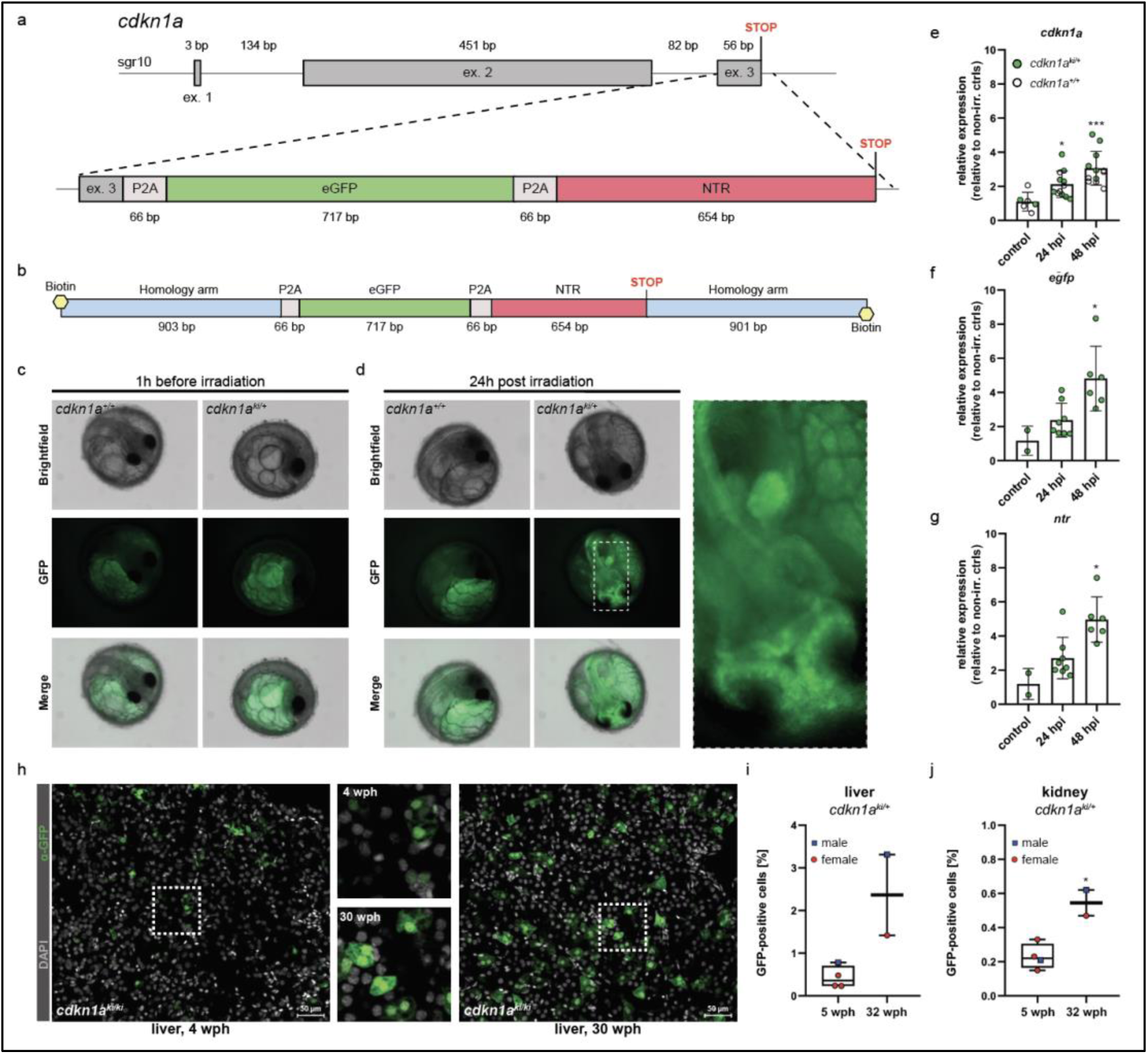
CRISPR/Cas9-mediated insertion of a reporter construct. **a**, The *cdkn1a* locus in *klara* was targeted via CRISPR/Cas9 to insert a reporter construct consisting of an eGFP and an NTR cassette, separated by P2A self-cleavage sites, allowing the detection and ablation of senescent cells. **b**, A 5’-biotinylated, double-stranded donor template flanked by two ~0.9 kb flanking arms was used for the target-specific insertion. **c**,**d**, Function of the reporter construct was tested by exposing embryos (*cdkn1a*^*+/+*^ and *cdkn1a*^*ki/+*^) to a γ-irradiation dose of 10 Gy. Representative images taken before irradiation only showed an autofluorescent signal originating from the yolk, which was observed in embryos of both genotypes **(c)**. 24 hours post irradiation (hpi), the presence of GFP-positive cells was detected only in *cdkn1a*^*ki/+*^ embryos **(d). e**, Gene expression analysis of *cdkn1a* in embryos (*cdkn1a*^*+/+*^ and *cdkn1a*^*ki/+*^) at 24 (n= 12) and 48 hpi (n= 12) compared to non-irradiated controls (n= 6). Expression of *cdkn1a* increased significantly (p_24hpi_: 0.011, p_48hpi_: 0.0004). **f**,**g**, For the expression of *egfp* **(f)** and *ntr* **(g)** at 24 (n= 8) and 48 hpi (n= 6) compared to non-irradiated controls (n= 2), *cdkn1a*^*+/+*^ embryos were excluded due to the absence of the reporter construct. Expressions of *egfp* and *ntr* were significantly increased at 48 hpi (p_*egfp*_: 0.045, p_*ntr*_: 0.006) compared to controls. For all qRT-PCRs, *rpl13a* was used as housekeeping gene. Relative gene expression was calculated using the ΔΔCT method. Student’s t-tests were computed to determine significant changes in gene expression. **h**, Detection of GFP-positive cells in the liver of a young (4 wph) and an old (30 wph) *cdkn1a*^*ki/ki*^ fish. A higher number of GFP^+^ cells was detected in the old liver sample. Images are displayed as extended depth of focus projections. In the young liver tissue, small individual GFP-positive cells were detected, whereas in the old liver a high prevalence of enlarged GFP-positive cells was identified (see zoom-in). **i**, Analysis of liver samples from young (5 wph, n= 4) and old (32 wph, n= 2) *cdkn1a*^*ki/+*^ fish via flow cytometry. **j**, In the kidney of *cdkn1a*^*ki/+*^ fish, a significantly higher proportion of GFP-positive cells (p: 0.012) was detected in old (n= 2) versus young (n= 4) animals using flow cytometry. Student’s t-test was computed to analyze for differences in cell number.

After the initial characterization of the *cdkn1a*^*ki/+*^ embryos, we raised them to adulthood and generated F_2_ offspring. Subsequently, we used immunofluorescence and flow cytometry to detect and quantify the occurrence of GFP-positive cells and their potential accumulation upon aging. As organs for analysis we selected liver and kidney from young and old *cdkn1a*^*ki/+*^ and *cdkn1a*^*ki/ki*^ animals. Staining for GFP revealed many more GFP^+^ cells in the liver of a 30 weeks-old *cdkn1a*^*ki/ki*^ fish than in a 4 weeks-old animal (Fig. 5h). Of note, GFP^+^ cells in older tissue were larger and often appeared clustered as opposed to smaller and scattered GFP^+^ cells in young tissue. Using FACS, we detected 0.4 and 0.2% GFP^+^ cells in the liver and kidney of 5 weeks-old *cdkn1a*^*ki/+*^ fish, respectively. This numbers increased to 2.4 and 0.5% in the liver and kidney of 32 weeks-old fish (Fig. 5i,j). These data suggest that the reporter that we have generated can be used to monitor *cdkn1a (p21)-*positive cells *in vivo* and that these cells accumulate upon aging.

## Discussion

Here, we report the generation of a transparent killifish line that is lacking melanophores, iridophores and xanthophores. Based on published literature, we have selected the three genes *mitfa, ltk* and *csf1ra* as targets for CRISPR/Cas9-mediated inactivation in *Nothobranchius furzeri* ^29,37,38^. We employed a single injection with three single guide RNAs that had been pre-selected and characterized. The observation that some of the injected F_0_ embryos showed a complete loss of melanophores demonstrated that the CRISPR/Cas9 system acts very efficiently in *N. furzeri*. This has been observed before ^22^ and is most likely explained by the long duration of the one-cell stage in this species of 2-3 hours during which Cas9 can act ^39^. Cas9 efficiency was further confirmed by the fact that for all three genes both alleles were inactivated in the F_0_ animals, many of which showed an almost complete transparency. This efficiency is comparable to the one reported for zebrafish, whereby a codon-optimized Cas9 protein was employed to target a reporter transgene and four endogenous loci. In this case, mutagenesis rates reached 75–99% ^40^.

The animals did not show any anomalies regarding phenotype and behavior and could be bred to homozygosity and kept as a stable line. While *klara* animals were initially fully transparent, male animals developed black pigments, particularly on fin appendages, which increased during their lifespan. This suggests that there is a second, *mitfa*-independent population of melanophores in killifish that appears at later life. In zebrafish it has been shown that the paralogous gene, *mitfb* might fulfill this role by activating *tyrosinase* expression ^41^. Whether or not this is also the case in killifish and whether *mitfb* might also be responsible for the remaining pigmentation in the retina remains to be determined. By inactivating *slc45a2* in *klara* fish, we obtained animals that were fully transparent regarding their body and the eye. This quadruple mutant might be particularly interesting for research on eye and retina regeneration and shows that *klara* animals can be used as a background for further gene inactivation.

We used the *klara* line for addressing mate choice in killifish. While breeding behavior was normal among *klara* animals, both wild type and *klara* animals preferred pigmented mating partners in competitive breeding situations. Here, mate choice for pigmented partners was more pronounced in females (>90%) than in males (approx. 75%). This might be explained by the fact that the difference in appearance of wild type and *klara* males is much more distinct than between the respective females. The choice for pigmented males might also be influenced by the larger weight of wild type males. Our observation is in line with an earlier report on the two-spotted gobies. In this case, males preferred to mate with more colourful females, which have bright yellow-orange bellies during the breeding season ^42^. It is very surprising that for mate choice *N. furzeri* seems to rely on visible cues, as their natural habitat is turbid with limited visibility ^43^. It is tempting to speculate that mate choice in *N. furzeri* could also be influenced by other traits like chemical signals including pheromones.

The main motivation to generate a transparent killifish line was the possibility to perform longitudinal studies regarding aging and regeneration and to be able to observe respective processes in real-time without having to sacrifice cohorts of animals at distinct time points. One of the hallmarks of aging is the accumulation of senescent cells that are characterized by the expression of specific markers including *cdkn1a (p21)* and *cdkn2a (p16)* ^33^. It is still a matter of debate whether senescent cells limit or extend lifespan ^44,45^. To visualize and address the role of senescent cells, we have integrated a cassette encompassing a fluorescence reporter as well as a nitroreductase allele into the *cdkn1a* locus of *klara* animals. The respective HDR template was flanked by 0.9 kb homology arms and carried biotinylated 5’-ends, as those modifications of double-stranded donor templates have been reported to increase HDR efficiency ^35,36^. Out of 35 injected embryos, four showed proper integration of the construct, corresponding to 11%. At least three F_0_ animals passed on the engineered allele to the next generation. Our analysis of *klara* embryos harboring the *GFP* allele in the *cdkn1a* locus had shown that the integrated reporter is functional and can be activated upon γ-irradiation. The characterization of respective adult animals demonstrated an accumulation of GFP^+^ cells with age, as determined by flow cytometry and immunofluorescence. Whether those cells are truly senescent cells remains to be determined. Since the reporter also encompasses a *ntr* allele, it shall be possible to delete the GFP^+^ and thus presumably senescent cells by administration of Metronidazole, a prodrug that is converted to a cytotoxic agent by NTR. With the transparent senescence reporter line, it will be possible to further characterize the function of senescent cells in development, aging and regeneration. The generation of additional reporters into different loci, e.g. *cdkn2a/b* or other senescent markers will also allow to address the possible heterogeneity of senescent cells ^46^.

We consider the *klara* fish that we describe here a valuable and versatile tool for research on aging, regeneration and behavior. This fish line will also be beneficial for colleagues interested in cancer biology and ecology. Beyond its potential to be used for the investigation of questions in biology *klara* animals can also contribute to the reduction of animal numbers, an aspect that gains increasing importance in biomedical research.

## Materials and Methods

### Fish husbandry

All the work reported here was performed in the wild type *N. furzeri* strain MZCS-08/122, which is originally derived from southern Mozambique ^27^, or the *klara* line. Fish are kept in single-housing at 26°C on a light:dark cycle of 12 hours each. Adult fish are fed once a day *ad libitum* with red mosquito larvae, whereas juvenile fish (up to 5 weeks post hatching) are fed with artemia twice a day. To obtain a high number of fertilized wild type oocytes for injections, multiple breeding groups consisting of ten fish (2 males, 8 females) were set up in 40 liter tanks. In order to obtain oocytes from *klara* fish, breeding groups of 1 male and 3 female were set up in tanks containing approximately 8.5 l. The sand box, which is necessary for the deposition of eggs, was always removed two days before and put back into the tank two hours before the injection. The eggs were collected with a sieve and were then used for microinjections. The routinely collection of eggs for line maintenance was done on a weekly basis. Eggs were put on coconut coir plates and stored at 29°C.

All fish were maintained in the Nothobranchius facility of the Leibniz Institute on Aging – Fritz Lipmann Institute Jena according to the German Animal Welfare Law. The performed experiments reported here were covered by the animal license FLI-17-016, FLI-20-001 and FLI-20-102, which were approved by the local authorities (Thüringer Landesamt für Verbraucherschutz).

### Design and synthesis of single-guide RNAs (sgRNAs)

Single-guide RNAs were designed based on the genome sequence provided by the *Nothobranchius furzeri* Genome Browser ^24^. Target sequences for sgRNAs had a length of 20 nucleotides followed by the PAM sequence-*NGG*. Only sequences containing a restriction site directly upstream of the PAM sequences were selected. A TAGG-overhang was added to the 5’-end of the forward sgRNA oligonucleotide and an AAAC-overhang to the 5’-end of the reverse complementary oligonucleotide (sg_*mitfa*_1: 5’-TAGG-TGAAATGGATTTCCTGATGG-3’ sg_*mitfa*_2: 5’-AAAC-CCATCAGGAAATCCATTTCA-3’, sg_*ltk*_1: 5’-TAGG-AACATCAAAAGGGAATTCAC-3’ sg_*ltk*_2: 5’-AAAC-GTGAATTCCCTTTTGATGTT-3’, sg_*csf1ra*_1: 5’-TAGG-CAGAGACACTTTTTCCATGG-3’ sg_*csf1ra*_2: 5’-AAAC-CCATGGAAAAAGTGTCTCTG-3’, sg_*slc45a2*_1: 5’-TAGG-TGACTACTGCCGCTCACAGT-3’ sg_*slc45a2*_2: 5’-AAAC-ACTGTGAGCGGCAGTAGTCA-3’). Complementary sgRNA oligonucleotides were annealed by heating them up to 95°C followed by gradually cooling by 1°C per 30 seconds. The annealed oligonucleotides were ligated into the *BsaI*-linearized pDR274 vector (Addgene, plasmid #42250). After the ligation, this vector, containing the sgRNA sequence, was transformed into *E. coli* TOP10 cells. Isolated plasmids were checked via sequencing for the correct presence of the sgRNA sequence. Using the *DraI* restriction enzyme a fragment of approximately 300 bp was excised from the plasmid containing the sgRNA and the T7 promoter sequence. This fragment was used as a template for the *in vitro* transcription, which was performed according to the manufacturer’s protocol of the mMESSAGE mMACHINE™ T7 Transcription Kit (Thermo Fisher Scientific Inc.). Quality of *in vitro* transcribed sgRNAs was controlled via RNA agarose gel electrophoresis.

### Design and synthesis of DNA donor templates for HDR

The assembly of the donor template for the insertion of a *P2A-eGFP-P2A-NTR* cassette into the *cdkn1a* locus of *klara* was done using the *NEBuilder*^*®*^ *HiFi DNA Assembly Cloning Kit. P2A* sites were added to the *eGFP* (derived from the *tol2* kit plasmid #395) and the *NTR* sequence (obtained from plasmid: *Myl7-LoxP-myctagBFP-LoxPNTRmCherry* ^47^) via PCR. The *NEBuilder Assembly Tool* was used for the design of oligonucleotides containing overlap sequences, which are required for the assembly of individual PCR fragments. 0.05 pmol of each fragment (flanking arms, *P2A-eGFP, P2A-NTR*, pGGC (pUC57-BsaI) backbone vector ^48^) were used together with 10 μl of the *NEBuilder*^*®*^ *HiFi DNA Assembly Master Mix*. The NEBuilder assembly reaction was performed for one hour at 50°C. 5 µl of this reaction were used for a subsequent transformation into chemically competent TOP10 *E. coli*. Isolated plasmids were checked via sequencing for correct template assembly. 5’-biotinylated oligonucleotides (bio_*cdkn1a*_fw: 5’-TCTTACACCAAACACCACAA-3’ bio_*cdkn1a*_rv: 5’-TAAAACATGCAGGATACCGG-3’) were used to amplify the template from the isolated and then linearized plasmid. The amplicon with the expected size was excised from the agarose gel and purified using the *NucleoSpin Gel and PCR clean-up* kit (Macherey-Nagel). In order to induce a DNA double-strand break in close proximity to the site of insertion, the following oligonucleotides for sgRNA synthesis were used: sg_*cdkn1a*_1: 5’-TAGG-AATATCACTCCCCGGATTTC-3’ sg_*cdkn1a*_2: 5’-AAAC-GAAATCCGGGGAGTGATATT-3’. Synthesis of this sgRNA was done as described above.

### Microinjections into *N. furzeri* oocytes

For microinjections, an injection mold (manufactured by GT-Labortechnik, Arnstein, Germany) forming single slots for the size of an *N. furzeri* embryo was used. Injection plates were freshly prepared by dissolving 1.5 g of agarose in 50 ml of 0.3 x Danieau’s medium. This solution was boiled in a microwave oven and then poured into a petri dish (94 mm x 16 mm). The injection mold was put in while the solution is still liquid and as soon as the agarose is hardened (after approximately 30 min), the stamp can be removed and the plate is ready to use. Fertilized embryos were lined up individually in each slot in a way that the cell was facing towards the direction of the injection needle. 0.3 x Danieau’s medium was added onto the plate until the eggs were completely covered. The injection was performed under a stereomicroscope using glass capillary needles, a pressure injector (World Precision Instruments) and a micromanipulator (Saur). The injection solution for the inactivation of target genes contained the sgRNAs (30 ng/µl each), *Cas9* mRNA (300 ng/µl), *GFP* mRNA (200 ng/µl) and phenol red. For knock-in approaches, the concentration of sgRNA and Cas9 mRNA was kept the same, whereas *GFP* mRNA was reduced to 100 ng/µl and 20 ng/µl of the HDR template were added. After injections, the embryos remained on the plate and were stored at 29°C until the next day. Using a fluorescence microscope, the injected embryos were sorted into GFP-positive and GFP-negative embryos. To reduce the risk of cross contaminations, GFP-positive embryos were transferred into single wells of a 96-well plate containing 0.3 x Danieau’s medium.

### DNA sampling

DNA samples were obtained either from caudal fin biopsies or whole embryos. For fin biopsies, a small part of the caudal fin was cut off from the anesthetized fish. For the extraction of DNA from whole embryos, the embryos were mechanically disrupted using a pipet tip. For fin biopsy samples 100 µl and for embryos 50 µl of NaOH (50 mM) was added to the sample and subsequently incubated at 95°C for 45 min. Afterwards, 10 µl or 5 µl respectively of Tris-HCl (1 M, pH 8.0) were added.

### Restriction enzyme digest

The sgRNA design enabled the use of restriction enzymes to check for the presence of mutations in the targeted genes. The region flanking the potential mutation site was amplified via PCR using the following oligonucleotides: *mitfa*_fw: 5’-TGCTTCACATACGTTTGCAG-3’ *mitfa*_rv: 5’-CAAAGGTCTGAGGGCTTTCC-3’, *ltk*_fw: 5’-TGTTCTGTCACCACCCTTGT-3’ *ltk*_rv: 5’-ACACTGCTATTACCAGGTTTGAC-3’, *csf1ra*_fw: 5’-CATAGATACCGTGCAAGCCTG-3’ *csf1ra*_rv: 5’-AGCCCAGGTATGAAATCCGT-3’, *slc45a2*_fw: 5’-GGATTTGGTGTTTTGGCCCT-3’ *slc45a2*_rv: 5’-GTAACTCGGCTCTAATCGTGC-3’. For the restriction enzyme digest, 20 µl of the PCR reaction were incubated over night at 37°C after adding 6.75 µl of ddH_2_O, 0.25 µl of the corresponding enzyme and 3 µl of the respective enzyme buffer (*mitfa*: *EcoRI, ltk*: *EcoNI, csf1ra*: *NcoI-HF, slc45a2*: *HypCH4III*). Samples from the control digest were analyzed on a 1% agarose gel. In the presence of a mutation, the restriction enzyme was not able to cleave the PCR fragment, whereas non-mutated sequences were still cleaved. As positive control, a PCR amplicon from a wild type fish was always included in order to verify that the restriction enzyme digest worked properly.

### High-resolution melting analysis (HRMA)

As soon as the exact mutations in the targeted gene loci were identified (from F_2_ generation on), genotyping was performed via HRMA. For this analysis, the CFX384™ Real-time PCR Detection System (Bio-Rad) was used. DNA samples obtained from either fin biopsies or whole embryo lysates were diluted 1:50 with H_2_O before use. The HRM analysis was done according to the protocol provided by the *Precision Melt Supermix* Kit (Bio-Rad; Catalog #172-5112). The following oligonucleotides were used for the HRM analysis: HRMA_*mitfa*_fw: 5’-CCTCACGAGTCTCTCTATCA-3’ HRMA_*mitfa*_rv: 5’-GCCCCATGAACCCAATATAA-3’, HRMA_*ltk*_fw: 5’-CCACAGACTCTTCCAGAAAT-3’ HRMA_*ltk*_rv: 5’-CTGATTATGAGGTGCGACTA-3’, HRMA_*csf1ra*_fw: 5’-AGTGTGTGGCTTTCAATTTG-3’ HRMA_*csf1ra*_rv: 5’-TTTCTGGTGAGTGTTTGTTA-3’. For the assessment of the genotypes, melt curves were analyzed using the *Precision Melt Analysis*^*TM*^ software (Bio-Rad).

### Isolation of nucleic acids

RNA isolation from fish tissues was done according to the manufacturer’s protocol of the RNeasy Mini Kit (Qiagen). Tissue homogenization using ceramic beats was performed with the TissueLyser II (2 min at 30 Hz). The optional on-column DNase digestion step was included. 20 µl of DEPC H_2_O were used for final elution. RNA isolation from FACS sorted cells was done according to the protocol of the *MagMaxTM-96 Total RNA Isolation Kit*. Isolation of RNA and DNA from whole embryos was done via phenol-chloroform extraction. The chorion of the embryos was mechanically disrupted using a pipet tip before 500 µl of TRIzol were added. Homogenization of the embryos was performed with ceramic beats using the TissueLyser II (2 min at 30 Hz). After an incubation at RT for 5 min, 200 µl of chloroform were added. Samples were then mixed for 15 s, incubated at RT for 3 min and then centrifuged at 12,000 x g for 20 min (4°C). The upper, aqueous phase, which contains the RNA, was transferred into a fresh tube and 1.1 volumes of isopropanol, 0.16 volumes of NaAc (2M, pH 4.0) and 1 µl of GlycoBlue were added. Samples were incubated at RT for 10 min and then centrifuged at 12,000 x g for 20 min (4°C). The supernatant was removed and the pellet was washed with 1 ml of 80% EtOH and centrifuged at 7,500 x g for 10 min (4°C). The supernatant was discarded and the pellet was air-dried. RNA pellet was dissolved in 20 µl of DEPC-H_2_O and stored at −80°C. The inter- and organic phase were used for extraction of DNA (required for genotyping PCR). 300 µl of EtOH (100%) were added, samples were incubated at RT for 2-3 min and then centrifuged at 2,000 x g for 5 min (4°C). Supernatant was discarded and pellet was incubated in 1 ml of 0.1 M sodium citrate in 10% EtOH for 30 min before centrifugation at 2,000 x g for 5 min (4°C). Supernatant was discarded and 1 ml of EtOH (75%) was added for 15 min before centrifugation at 2,000 x g for 5 min (4°C). Pellet was air-dried and afterwards dissolved in 15 µl of NaOH (8 mM).

### cDNA synthesis and gene expression analysis

For cDNA synthesis, 500 ng of RNA from fish tissues and whole embryos or 25 ng of RNA from FACS sorted cells were used. cDNA synthesis was performed according to the instructions of the *iScript cDNA Synthesis Kit* (Bio-Rad). qRT-PCRs were performed in 384-well plates using 2x SYBR Green Mix and the CFX384 Real-Time System (Bio-Rad). Reaction mix included 3 µl of cDNA (diluted 1:5 in DEPC H_2_O), 0.4 µl of each oligonucleotide, 1.2 µl DEPC H_2_O and 5 µl of 2x SYBR Green Mix. Gene expression levels were determined using the following oligonucleotides: q_*mitfa*_fw: 5’-TGAAGCAAGTACTGGACAAG-3’ q_*mitfa*_rv: 5’-TCCAGTAGAGTCAGAAGTCC-3’, q_*ltk*_fw: 5’-CTGGGAGGAATCCGCTTA-3’ q_*ltk*_rv: 5’-AGTGAGACCAGTGCAGAG-3’, q_*csf1ra*_fw: 5’-AGTTCAAATGTATCAGAGACCT-3’ q_*csf1ra*_rv: 5’-TATCCTGCTCCGAGAATCAT-3’, q_*gfp*_fw: 5’-AAGGGCATCGACTTCAAGGA-3’ q_*gfp*_rv: 5’-GGCGGATCTTGAAGTTCACC-3’, q_*ntr*_fw: 5’-CTTTTGATGCCAGCAAGAAA-3’ q_*ntr*_rv: 5’-GAAGCCACAATAAAATGCCA-3’, q_*cdkn1a*_fw: 5’-ATGTGCAGAGGGATGGCTAC-3’ q_*cdkn1a*_rv: 5’-CCTCCAGATCTTTACGCAG-3’. For normalization the housekeeping gene *rpl13a* (q_*rpl13a*_fw: 5’-ACTGTCAGAGGCATGCTTCC-3’ q_*rpl13a*_rv: 5’-TGCTCTGAAAATTGTGCGCC-3’) was used.

### Whole kidney marrow analysis via flow cytometry

Kidneys were dissected from fish and immediately pressed through a 40 μm cell strainer (placed on top of a 50 ml falcon) using a syringe plunger. Strainer and plunger were rinsed with 1 ml of PBS each. Cells were pelleted by centrifugation at 330 x g for 5 min (4°C). Supernatant was discarded and the cell pellet was dissolved in 300 μl of PBS. Analysis of the WKM was done using a BD FACS AriaTM IIIu. The gating strategy was chosen as described for the analysis of the WKM of zebrafish ^32^.

### Irradiation of *N. furzeri* eggs

For γ–irradiation of F_1_ eggs from the *cdkn1a*-reporter line, eggs were placed into 12-well plates (one egg per well) containing 1 ml of 0.3x Danieau’s medium. Eggs were irradiated in the 12-well plate with a dose of 10 Gy using a Gammacell^®^ 40 Exactor (Best Theratronics Ltd.). The presence of an eGFP signal was checked using an Axio Zoom.V16 with ApoTome (Zeiss).

### Genotyping of eggs and fish from the *cdkn1a*-reporter line

DNA was extracted from eggs or fin biopsies as described above. To check for the presence of the reporter construct in the *cdkn1a* locus, the following primer pair was used: *cdkn1a*_insertion_fw: 5’-TATTTCTCTGGTGTTTGCCT-3’ *egfp*_insertion_rv: 5’-TGATATAGACGTTGTGGCTG-3’. To discriminate between the three different genotypes, the following primer pair was used: *cdkn1a*_genotyping_fw: 5’-CTACAGATCCAGCGTCATC-3’ *cdkn1a*_genotyping_rv: 5’-CCAAGAGAACCAGACAAAGA-3’. For *cdkn1a*^*+/+*^ animals an amplicon with a size of 393 bp was expected, whereas a 1,893 bp fragment occurred in *cdkn1a*^*ki/ki*^ fish. In *cdkn1a*^*ki/+*^ animals both amplicons were present.

### Analysis of GFP-positive cells from the *cdkn1a*-reporter line

Organs (liver and kidney) were removed from the fish and put into 1x PBS on ice. Liver samples were then transferred into 1.5 ml tubes containing 900 µl of sterile PBS and 100 µl of Collagenase I. After an incubation for 1h at 32°C at 550 rpm, 100 µl of FBS were added and tubes were placed on ice. Livers were then pushed through a 100 µm cell strainer placed on a 50 ml tube using syringe plunger. Cell strainer and plunger were rinsed twice with 1 ml of PBS each. After centrifugation at 250 x g for 5 min (4°C), the pellet was resuspended in 1 ml PBS containing 25 µl of Collagenase I and 3 µl Dispase followed by an incubation for 30 min at 37°C. Afterwards, 100 µl of FBS were added and the solution was pushed through a 40 µm cell strainer placed on a 50 ml tube. Kidney samples were directly placed on a 40 µm cell strainer, 1 ml of PBS with 1% FBS was added and then pushed through the cell strainer using a syringe plunger. Cell strainer and plunger were rinsed twice with 1 ml of PBS each.

After centrifugation at 250 x g for 5 min (4°C), the pellet was resuspended in 200 µl PBS containing 1% FBS. The cell suspension was stained with 5 nM Sytox Red Dead Cell Stain (Invitrogen) to identify dead cells and were subsequently sorted on a BD FACS AriaIIIu Cell Sorter using a 100 µm Nozzle. After doublet and dead cell exclusion, the GFP-positive cells were sorted in PBS. The GFP+ gate was set according to the negative control obtained from respective wild type tissues. Data were analyzed using FlowJo v10 Software (BD).

### Immunofluorescence staining

Liver tissues were fixed overnight in 4% paraformaldehyde in PBS and subsequently washed three times in PBS-T (0.2% Tween in PBS). Tissues were placed for 5 min in 5% sucrose, then for 2h in 20% sucrose and subsequently overnight in 30% sucrose. Tissues were embedded and shock frozen in molds filled with NEG-50™ cryosection medium. The tissue was cut into 20 µm slices. After defrosting for 25 min, the sections were washed four times for 10 min at RT in PBS-T (0.2% Tween) and were permeabilized by a short washing step with permeabilization solution (0.1% Tween, 0.3% Triton X 100 in PBS). Immediately afterwards, the sections were washed again two times for 10 min at RT in PBS-T (0.2% Tween) and incubated in blocking buffer (2% BSA and 10% NGS in PBS-T (0.2% Tween)) for 1h at RT. Afterwards, the samples were incubated overnight at 4°C with an anti-GFP antibody (Thermo Fisher Scientific Inc., United States: A-11122, rabbit) diluted 1:200 in blocking buffer. Sections were then washed four times for 10 minutes at RT in PBS-T (0.2% Tween) prior incubation with bisbenzimide Hoechst 33258 and a secondary anti-rabbit Alexa Fluor^®^ 546 antibody (Thermo Fisher Scientific Inc., United States: A-11071, goat) diluted 1:500 in blocking solution for 1h at RT. After multiple washing steps in PBS-T (0.2% Tween), the slides were mounted with 70µl ProLong^®^ Diamond antifade reagent (Thermo Fisher Scientific Inc.). Before imaging, the samples were incubated at 4°C overnight. Image stacks were recorded as optical sections with the Axio Imager 2 equipped with an ApoTome.2 slider (Zeiss, Germany). The ZEN 3.4 software (Zeiss, Germany) was used to process the images and to create extended depth of focus projections of the acquired z-stacks.

### Statistical analysis

Data were analyzed depending on the experimental setup via t-test or One-Way ANOVA followed by Tukey’s post hoc test. Equal or unequal variance was determined via F-Test followed by a Student’s or Welch’s t-test, respectively. Significant changes are indicated by * if p≤0.05, ** if p≤0.01, and *** if p≤0.001.

## Acknowledgements

We thank Hanna Reuter for suggesting the name *klara*, Nils Hartmann for the movie on wild type killifish mating, Annekatrin Richter for providing the picture on oocyte injection and Hakar Aliyas, Caglar Avci, Michelle Burkhardt, Christina Ebert, Gabriele Günther, Maleen Hofmann, Erik Hüttenrauch and Dagmar Kruspe for technical support. We are very grateful to members of FLI’s killifish facility, most notably to Simone Dunkel, Martin Neumann, Marcus Schmidt, Uta Naumann and Beate Hoppe. We also would like to thank members of the Core Facility Flow Cytometry, namely Johanna Schleep, Simone Tänzer and Katrin Schubert for their contribution. This project was made possible by funding from the Carl Zeiss Foundation in the context of the IMPULS consortium (project number P2019-01-006) to C.E. and a fellowship from the Leibniz Graduate School on Ageing and Age-Related Diseases (LGSA) to J.K. The FLI is a member of the Leibniz Association and is financially supported by the Federal Government of Germany and the State of Thuringia.

## Author Contributions

J.K. and C.E. conceived the study. C.A. supported characterization of the senescence reporter fish. V.L.H. was instrumental in establishing HDR-based targeted insertions in the lab and performed the imaging of the senescence reporter line. J.K. performed all other experiments and data analyses. C.E. supervised the study. J.K. and C.E. wrote the manuscript.

## Competing Interests statement

The authors declare no competing interests.

---

**Extended Data Fig. 1:**
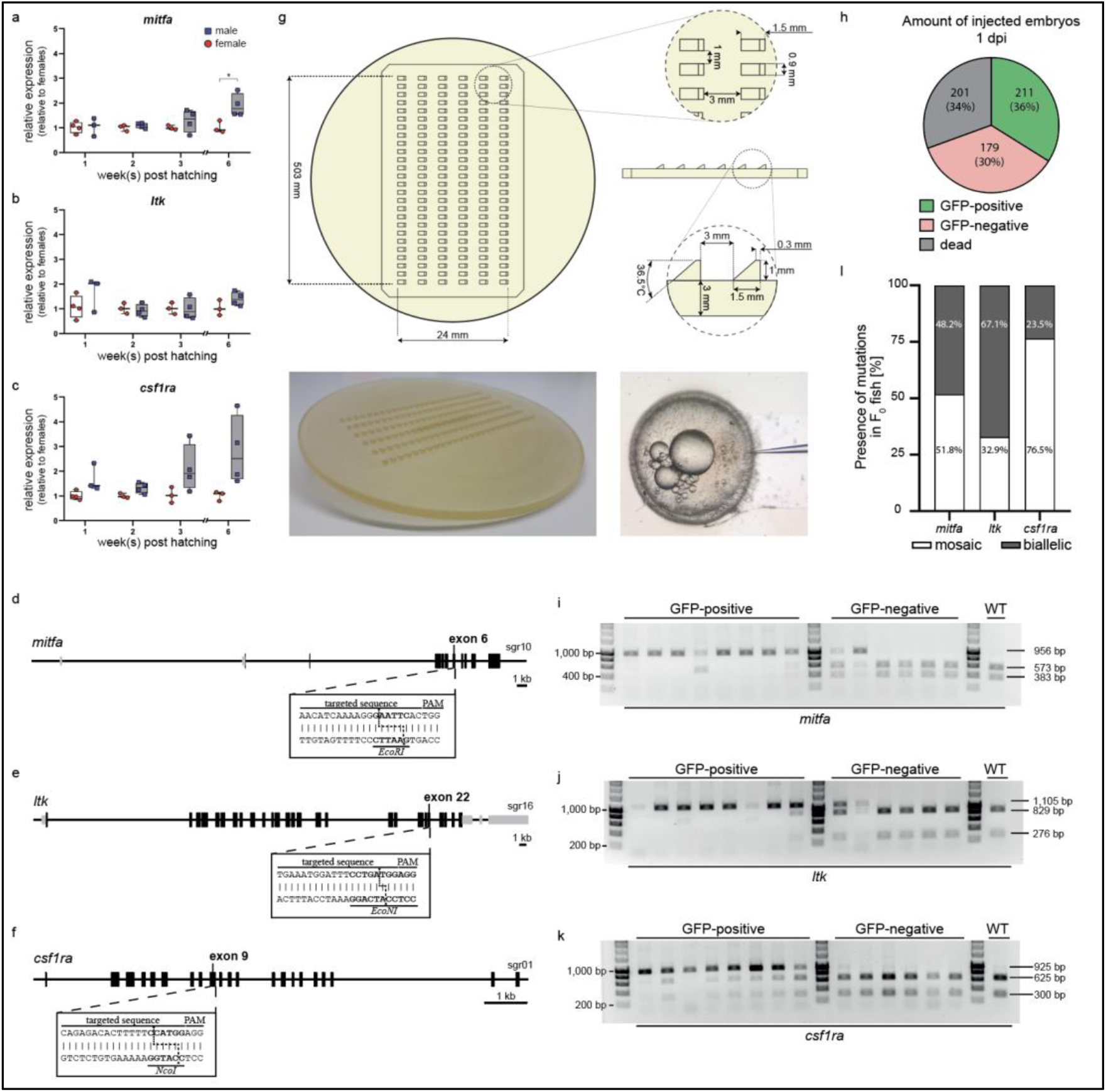
Analyzing and targeting the *mitfa, ltk* and *csf1ra* loci in *N. furzeri*. **a**,**b**,**c**, Intersexual comparison of the *mitfa* **(a)**, *ltk* **(b)** and *csf1ra* **(c)** expression in the skin of wild type fish at 1 (n_male_= 3, n_female_= 4), 2 (n_male_= 4, n_female_= 3), 3 (n_male_= 4, n_female_= 3) and 6 wph (n_male_= 4, n_female_= 3) using qRT-PCR. A significant change in gene expression between females and males was detected for *mitfa* at 6 wph (p: 0.031). Expression levels were normalized to the one in females at the respective age. *Rpl13a* was used as housekeeping gene. Relative gene expression was calculated using the ΔΔCT method. Student’s or Welch’s t-tests were computed to determine significant changes in gene expression. **d**,**e**,**f**, Schematic representation of the *mitfa* **(d)**, *ltk* **(e)** and *csf1ra* **(f)** loci in *N. furzeri*. Per gene one sgRNA was designed targeting a sequence part in exon 6 of *mitfa*, exon 22 of *ltk* and exon 9 of *csf1ra*. Recognition sites for restriction enzymes (*mitfa*: *EcoRI, ltk*: *EcoNI, csf1ra*: *NcoI*) directly upstream of the respective PAM sequence were used to check for the presence of mutations. **g**, Injection mold stamp designed for the preparation of plates for microinjections. Using this mold, single slots were generated on an agarose plate that stabilize and hold freshly fertilized *N. furzeri* eggs for microinjections (Picture showing oocyte injection was kindly provided by Annekatrin Richter, FLI Jena). **h**, One day after injection (dpi), injected eggs (n= 591) were screened for the presence of a GFP signal. 36% of injected eggs were GFP-positive (n= 211), 30% GFP-negative (n= 179) and 34% were dead (n= 201). **i**,**j**,**k**, Restriction enzyme digests to analyze the presence of mutations in *mitfa* **(i)**, *ltk* **(j)** and *csf1ra* **(k)** of 8 randomly selected GFP-positive, 6 randomly selected GFP-negative embryos and a wild type control. Non-cleaved products (*mitfa*: 956 bp, *ltk*: 1.105 bp, *csf1ra*: 926bp) indicated the presence of a mutation in the respective locus. Occurrence of two smaller products (*mitfa*: 383 bp + 573 bp, *ltk*: 276 bp + 829 bp, *csf1ra*: 300 bp + 626 bp) indicated the presence of the wild type sequence. **l**, Based on restriction enzyme digests, the occurrence of mutations in the *mitfa, ltk* and *csf1ra* locus of 85 F_0_ fish was analyzed. All fish (85/85) carried a mutation in *mitfa* (mosaic: 51.8%; biallelic: 48.2%), *ltk* (mosaic: 32.9%; biallelic: 67.1%) and *csf1ra* (mosaic: 76.5%; biallelic: 23.5%).

**Extended Data Fig. 2:**
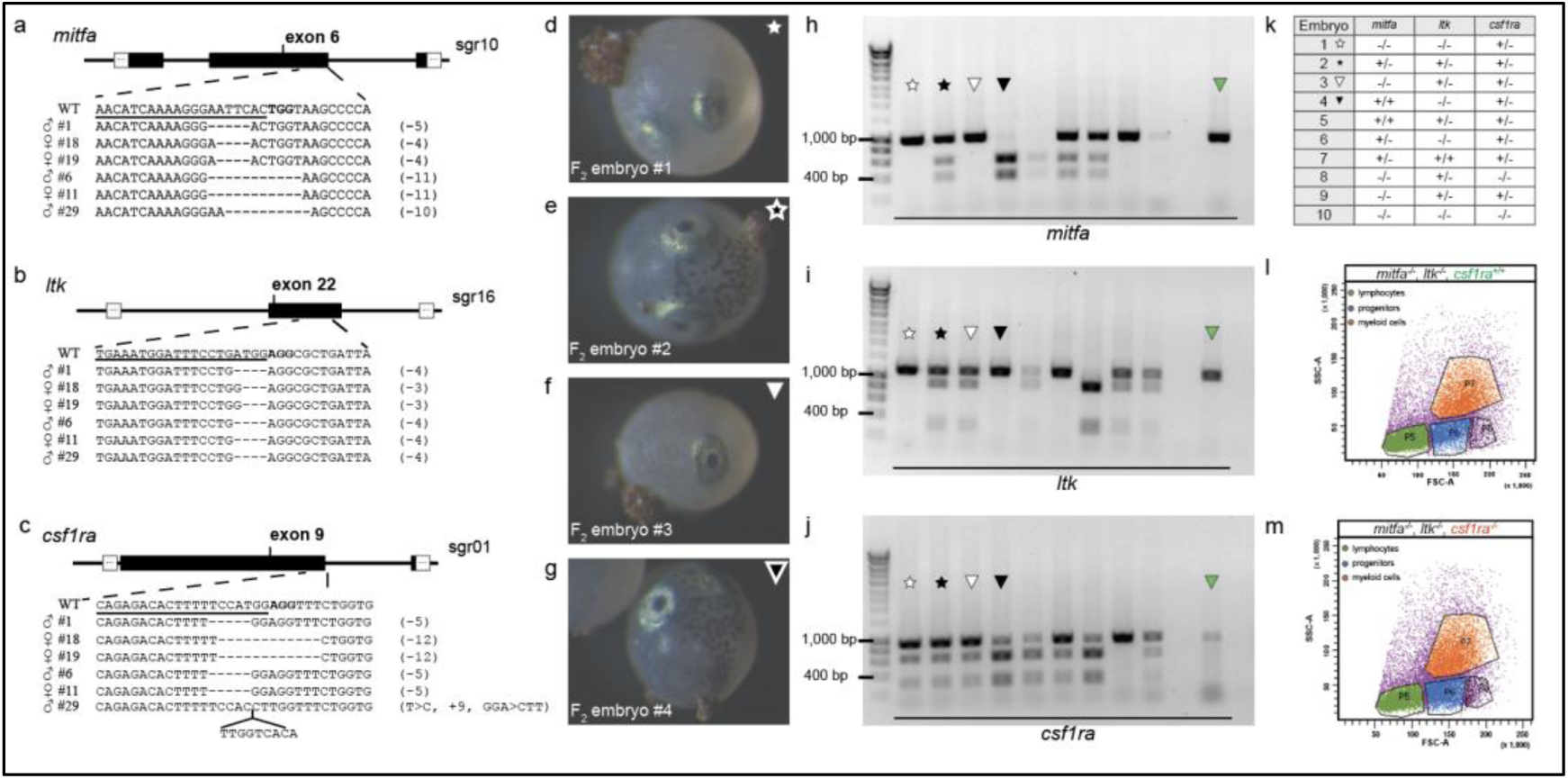
Characterization of *klara* fish. **a**,**b**,**c**, Via sequencing, indel mutations were identified in *mitfa* **(a)**, *ltk* **(b)** and *csf1ra* **(c)** of F_1_ fish, originating from an outcross of F_0_ animals with wild type fish. CLUSTALW was used for sequence alignments, in which each line represents one fish. Deletions are indicated by dashed lines. The sequence part targeted by the respective sgRNA is underlined. **d**,**e**,**f**,**g**, Phenotypic analysis of F_2_ embryos, resulting from an incross of two triple heterozygous (*mitfa*^*+/−*^,*ltk*^*+/−*^,*csf1ra*^*+/−*^) fish. The absence of melanophores was detected in embryos #1 **(d)** and #3 **(f)**. In contrast, melanophores were present in embryos #2 **(e)** and #4 **(g). h**,**i**,**j**, Restriction enzyme digests to analyze the presence of mutations in *mitfa* **(h)**, *ltk* **(i)** and *csf1ra* **(j)** of 10 randomly selected F_2_ embryos. Non-cleaved products (*mitfa*: 956 bp, *ltk*: 1.105 bp, *csf1ra*: 926bp) indicated the presence of a mutation in the respective locus. Occurrence of two smaller products (*mitfa*: 383 bp + 573 bp, *ltk*: 276 bp + 829 bp, *csf1ra*: 300 bp + 626 bp) indicated the presence of the wild type sequence. **k**, Assessment of genotypes based on the results from the restriction enzyme digests. Embryo #10 showed homozygous mutation in all three analyzed loci (*mitfa*^*-/-*^,*ltk*^*-/-*^,*csf1ra*^*-/-*^). **l**,**m**, Forward scatter (FSC-A) versus side scatter (SSC-A) plot of the whole kidney marrow (WKM) of a *mitfa*^*-/-*^,*ltk*^*-/-*^,*csf1ra*^*+/+*^ fish **(l)** and a *mitfa*^*-/-*^,*ltk*^*-/-*^,*csf1ra*^*-/-*^ fish **(m)**.

**Extended Data Fig. 3:**
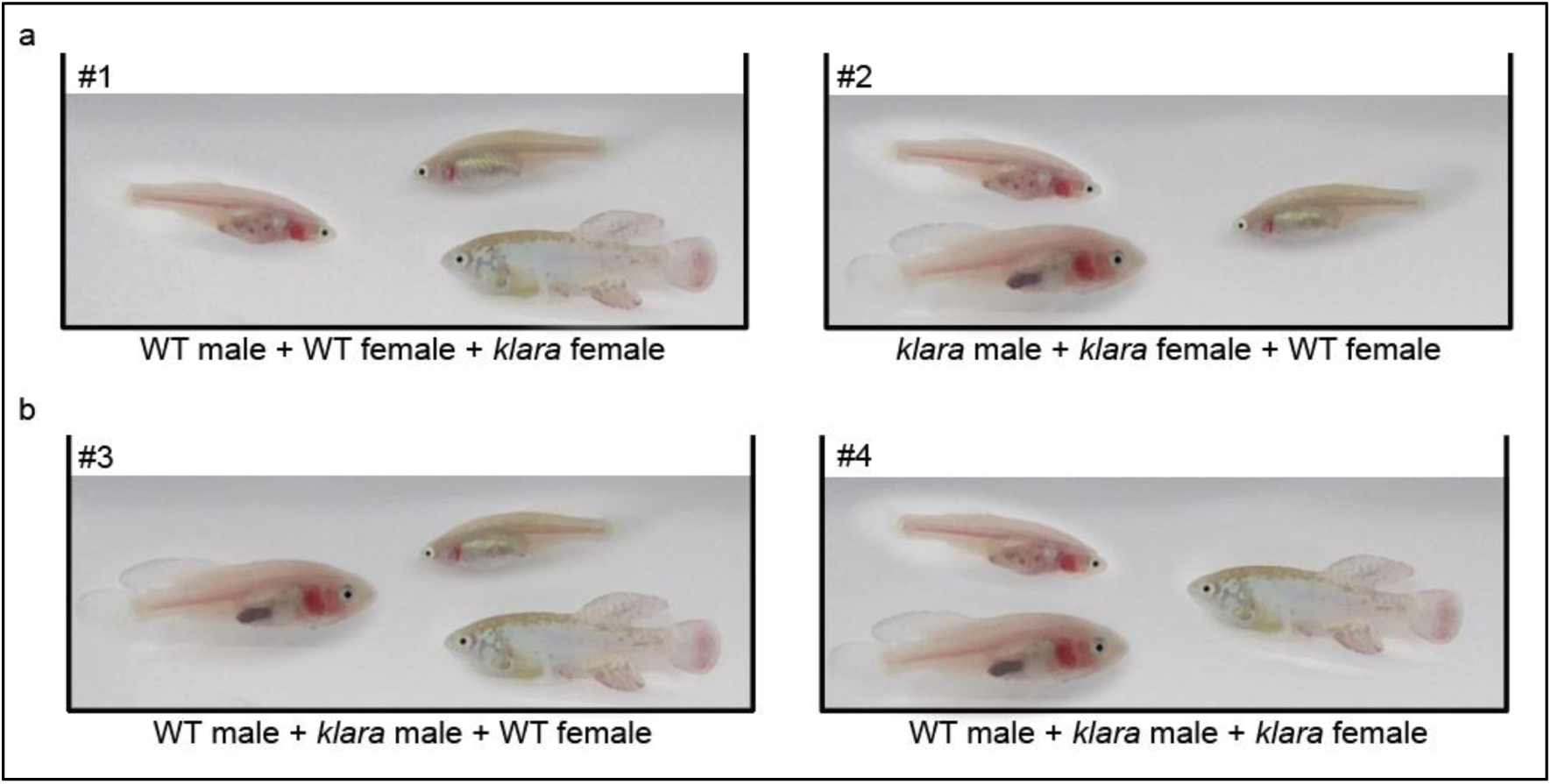
Groups for competitive breeding experiment. **a**, Breeding trios with two females (1x *klara* and 1x wild type) and either one wild type male (group #1) or one *klara* male (group #2). **b**, Composition of breeding trios with two males (1x *klara* and 1x wild type) and either one wild type female (group #3) or one *klara* female (group #4). Three tanks were analyzed per group.

**Extended Data Fig. 4:**
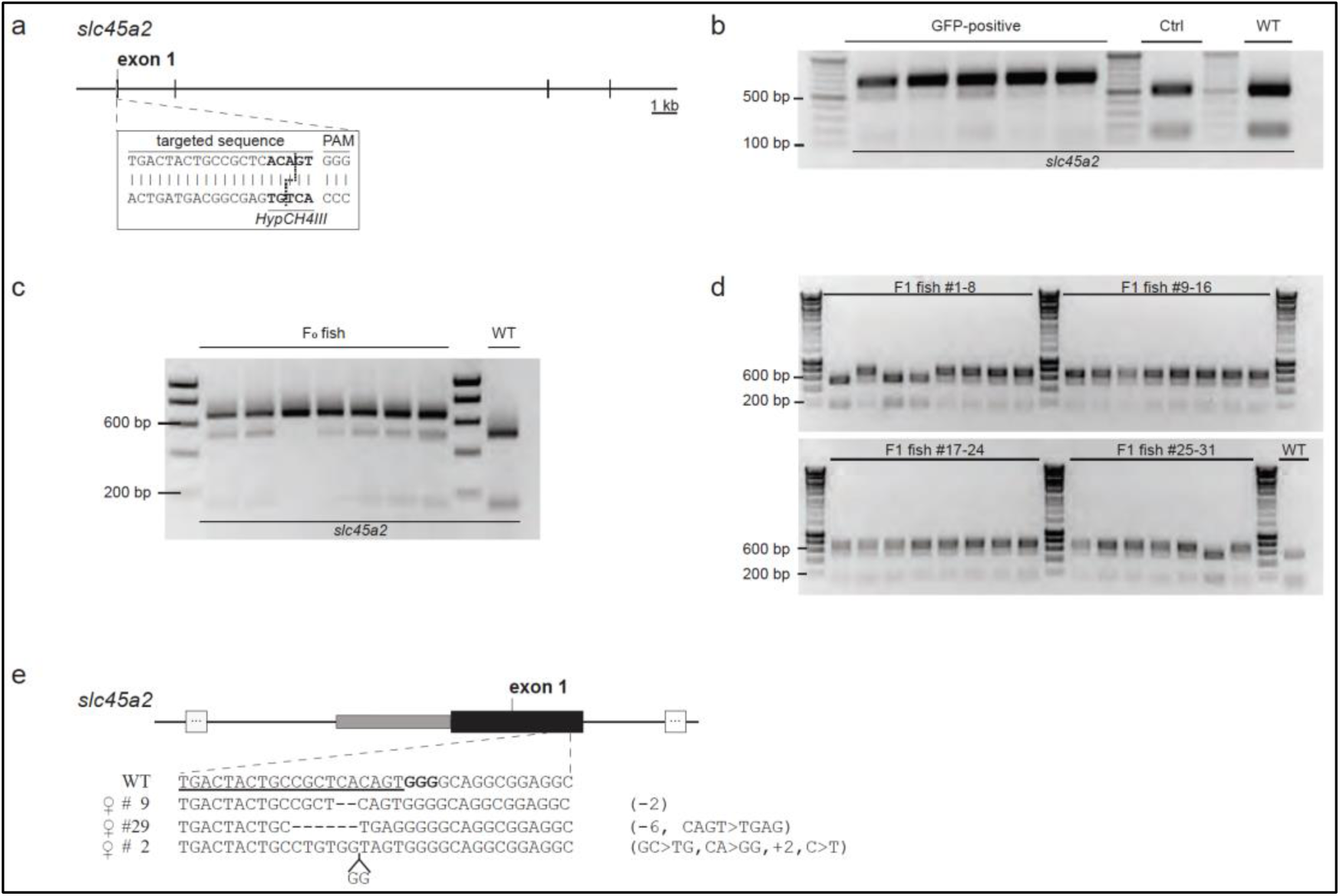
Inactivation of the *slc45a2* locus. **a**, Schematic representation of the *slc45a2* locus in *N. furzeri*. An sgRNA was designed targeting a sequence part in exon 1. The *HypCH4III* recognition site upstream of the PAM sequence allowed checking for the presence of mutations via restriction enzyme digest. **b**, All GFP-positive embryos (5/5) from microinjections of the *slc45a2* sgRNA into wild type oocytes showed the presence of a non-cleaved fragment (652 bp) indicating the presence of a mutation. Cleaved fragments (152 bp + 500 bp) were detected in uninjected and wild type control samples. **c**, Analyzing seven hatched F_0_ animals via restriction enzyme digest revealed the presence of mutations in all (7/7) fish indicated by the non-cleaved product. One fish (#3) did not show the additional cleaved fragments (152 bp + 500 bp) indicating a biallelic mutation in *slc45a2*. **d**, Mutations in *slc45a2* were present in 27/31 F_1_ fish originating from the outcross of an F_0_ animal with a wild type fish. **e**, Various indel mutations were identified in F_1_ fish via sequencing. CLUSTALW was used for sequence alignments, in which each line represents one fish. Deletions are indicated by dashed lines. The sequence part targeted by the used sgRNA is underlined.

**Extended Data Fig. 1:**
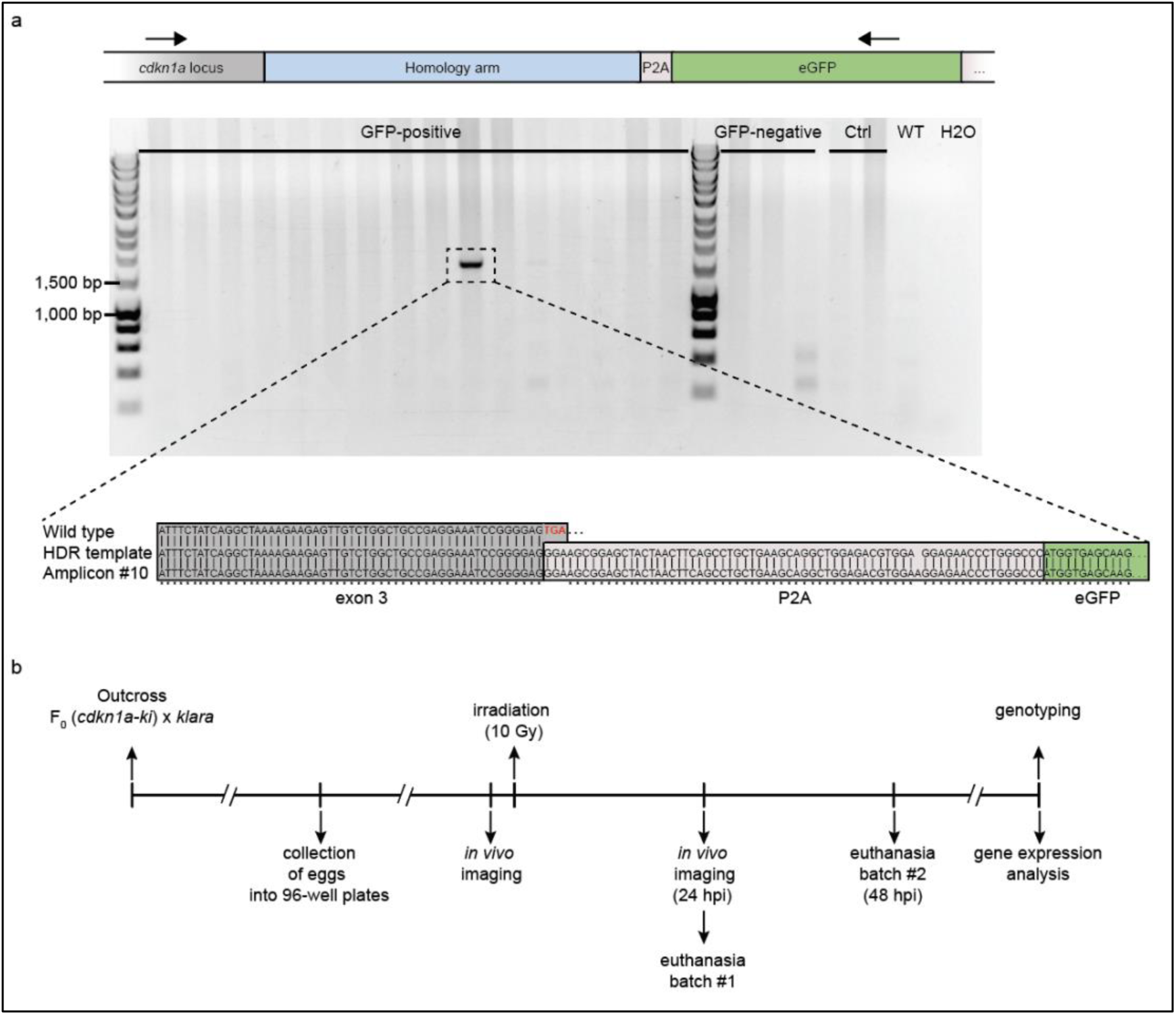
Analysis of embryos from the senescence reporter line. **a**, Using a primer pair combination in which one oligonucleotide binding site was located within the *cdkn1a* locus outside of the sequence that was covered by the homology arm of the donor template and the other one binding within the insert, 1 out of 16 randomly selected GFP-positive embryos showed an amplicon indicating the insertion of the construct. Proper insertion, i.e. lack of the endogenous STOP codon followed by the presence of the *P2A-eGFP-P2A-NTR* sequence, was verified via sequencing of the PCR amplicon. **b**, Workflow of the reporter construct test via γ-irradiation (10 Gy) in embryos (*cdkn1a*^*+/+*^ and *cdkn1a*^*ki/+*^).

## Notes

### Competing Interest Statement

The authors have declared no competing interest.

